# Pain hypersensitivity is dependent on autophagy protein Beclin 1 in males but not females

**DOI:** 10.1101/2023.06.13.544784

**Authors:** Theresa H. Tam, Wenbo Zhang, YuShan Tu, Janice L. Hicks, Sophia Farcas, Michael W. Salter

## Abstract

Chronic pain is a pervasive health, social, and economic problem affecting 1 in 5 individuals around the world. Increasingly, it is understood that alterations in fundamental cell biological processes are critical for chronic pain. A prominent cellular process is autophagy but whether it plays a role in pain is unknown. To investigate whether autophagy is involved in pain processing and is targetable for pain relief, we focused on Beclin 1, a component of the class III phosphatidylinositol 3-kinase (PI3K) complex necessary for initiating autophagy. Here, we found that inflammatory pain hypersensitivity in male mice lacking one allele of *Becn1* is significantly greater than that in wild type mice. By contrast, in female mice, loss of *Becn1* did not affect inflammation-induced pain hypersensitivity. Further, intrathecal delivery of an activator of Beclin 1, tat-beclin 1, reversed mechanical hypersensitivity induced by peripheral inflammation or peripheral nerve injury in males. Tat-beclin 1 also prevented mechanical hypersensitivity induced by exogenous brain-derived neurotrophic factor (BDNF), a core mediator of inflammatory and neuropathic pain in the spinal dorsal horn in males. Pain signaling pathways converge on enhancement of *N*-methyl-D-aspartate receptors (NMDARs) in spinal dorsal horn neurons. We found that loss of Beclin 1 increases expression of the pain-critical NMDAR subunit, GluN2B, in the dorsal horn and upregulates synaptic NMDAR-mediated currents in dorsal horn neurons from males but not females. From our converging lines of evidence, we conclude that inhibition of Beclin 1 in the dorsal horn is critical in mediating inflammatory and neuropathic pain signaling pathways in males. Our findings provide the basis for sex-specific therapeutic approaches targeting pain with a new class of analgesics - activators of Beclin 1.

## INTRODUCTION

Chronic pain is a global health problem, affecting approximately 20% of the population (*1, 2*). Still, current therapeutic options available are limited by poor efficacy and unacceptable side effects (*3, 4*). Physiological pain is critical for detecting and responding to tissue-damaging, or potentially tissue-damaging, stimuli. By contrast, pathological pain, such as inflammatory pain stemming from tissue inflammation or neuropathic pain from direct injury to or lesions of the nervous system, serves no known protective function (*5*). In pathological pain states, molecular changes occur within the dorsal horn of the spinal cord that lead to increased firing of dorsal horn nociceptive neurons resulting in enhanced transmission in pain pathways to the brain and pain hypersensitivity (*6*). Therefore, the spinal dorsal horn is a rich source for discovering signaling pathways and mechanisms that may be the basis for effective therapies for chronic pain.

An unappreciated pathway in the spinal dorsal horn is macroautophagy, herein referred to as autophagy. As a fundamental cell biological process, autophagy mediates degradation of proteins and other cellular components (*7*). At the core of autophagy is an intracellular cascade involving a series of protein complexes and intracellular organelles. A key protein complex, through which autophagy is initiated, is the class III phosphatidylinositol 3-kinase (PI3K). An essential component of the class III PI3K is the protein Beclin 1, which is assembled into the complex during autophagy initiation (*8, 9*). In the present study we explored the potential for autophagy to be involved in pain processing by experimentally manipulating Beclin 1.

## RESULTS

### Reduced Beclin 1 increases inflammatory pain hypersensitivity in male mice

We investigated the potential involvement of Beclin 1 in pain processing using mice in which expression of Beclin 1 is reduced through genomic deletion (*10, 11*). Homozygous deletion of *Becn1* is embryonically lethal (*10*) and therefore we used mice with monoallelic deletion of *Becn1* (*Becn1^+/-^*). As the dorsal horn of the spinal cord is the initial pain processing site of the central nervous system, we investigated the level of Beclin 1 protein in the spinal dorsal horn. We found in *Becn1^+/-^* mice that Beclin 1 protein level was statistically significantly less than that in *Becn1^+/+^* mice in males (Fig. 1A1) and in females (Fig. 1A2). Thus, in both sexes there is reduced expression of Beclin 1 in the spinal dorsal horn of *Becn1^+/-^* mice.

**Fig. 1.**
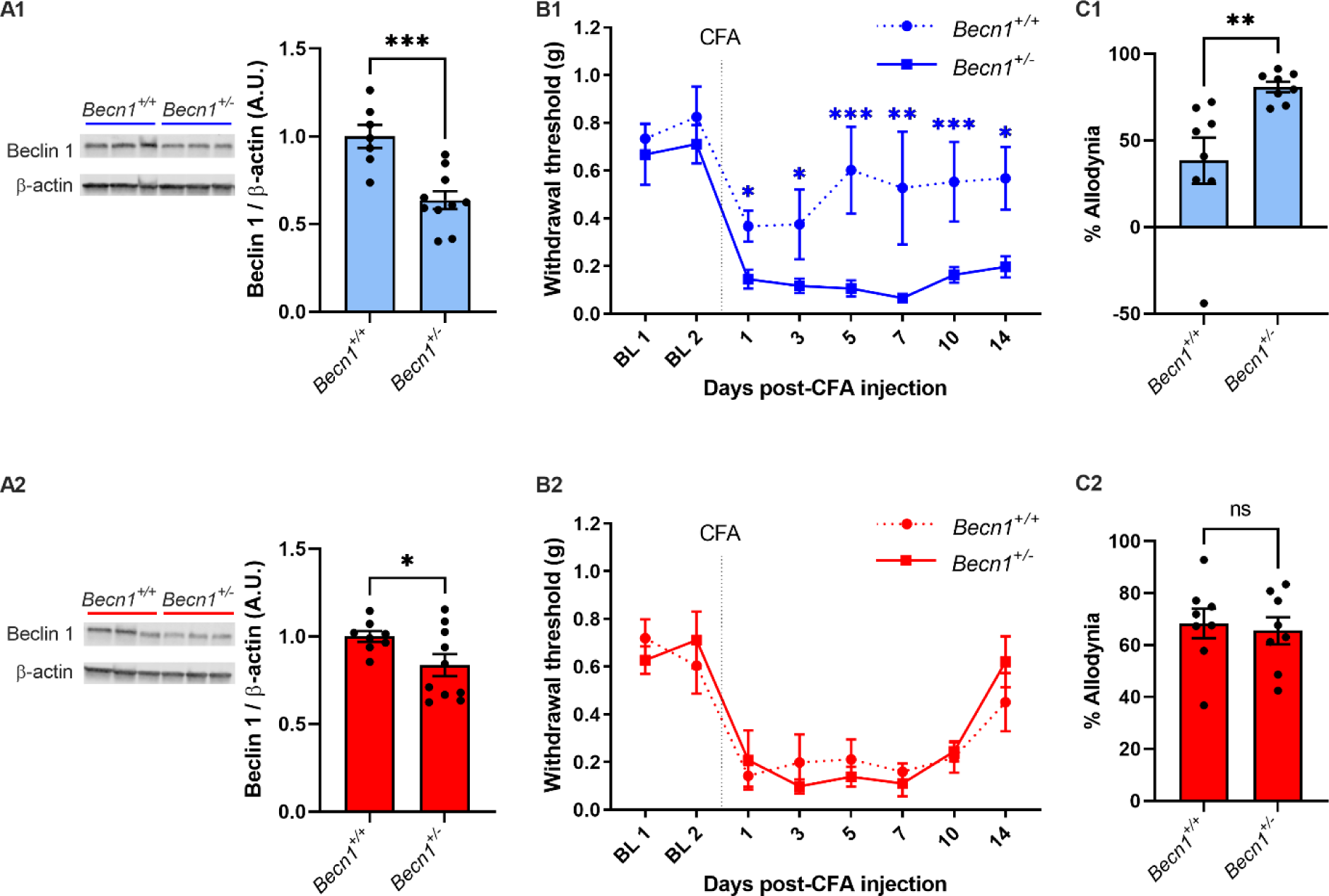
Monoallelic deletion of *Becn1* increases inflammation-induced pain hypersensitivity in males but not females. **(A)** Immunoblot of protein expression of Beclin 1 in spinal dorsal horn tissue in males *(A1)* and females *(A2)*. Males *Becn1^+/+^* n=7, males *Becn1^+/-^* n=10; females *Becn1^+/+^* n=8, females *Becn1^+/-^*n=10. *p<0.05, ***p<0.001, student’s t-test. **(B)** Mechanical paw withdrawal thresholds (PWT) of *Becn1^+/+^* or *Becn1^+/-^*males *(B1)* and females *(B2)* at two baseline (BL) measurements and multiple days following intraplantar CFA injection. **(C)** Quantification as percent allodynia of CFA (area over the curve function for PWT-time curve) in *Becn1^+/+^* or *Becn1^+/-^* males *(C1)* and females *(C2)*. n=8; *p<0.05, **p<0.01, ***p<0.001, \**Becn1^+/+^* vs *Becn1^+/-^*, Mann-Whitney U test.

A cardinal sign of pathological pain is hypersensitivity to innocuous mechanical stimuli, commonly referred to as allodynia (*12*). We assessed sensitivity to mechanical stimuli by applying nylon filaments of varying bending force to the hind paw of mice to determine the paw withdrawal threshold (PWT). In each sex, we found that, in naïve mice, the PWT was consistent at baseline (Fig. 1B) and that the PWT in *Becn1^+/-^* mice was indistinguishable from that in *Becn1^+/+^* mice (Fig. 1B). Thus, in mice with reduced Beclin 1 there was no alteration in basal mechanical pain threshold.

To investigate inflammatory pain hypersensitivity in *Becn1^+/-^*mice, we used intraplantar injection of Complete Freund’s Adjuvant (CFA), a well-established model of inflammatory pain (*13*), which causes a reduction in mechanical PWT indicative of mechanical hypersensitivity (Fig 1B). We found in males that PWT was reduced significantly more in *Becn1^+/-^* mice than in *Becn1^+/+^*mice, across the 14 day measurement period following CFA injection (Fig. 1B1; two-way ANOVA for genotype F(1,14)=5.8, p=0.03). Also, *Becn1^+/-^*male mice had significantly greater overall percentage allodynia than did male *Becn1^+/+^* mice, as assessed by comparing the area over the PWT-time curves (Fig. 1C1). By contrast, in females the CFA-induced reduction in PWT did not differ between the two genotypes (Fig. 1B2; two-way ANOVA for genotype F(1,14)=0.006, p=0.9), nor was percent allodynia in female *Becn1^+/-^* mice different from that in *Becn1^+/+^*(Fig. 1C2).

As paw withdrawal is a motor response, we investigated whether reducing Beclin 1 affects motor function. We used the accelerating rotarod test and found that the amount of time that *Becn1^+/-^* mice spent on the accelerating rotarod was not different from that spent by *Becn1^+/+^* mice (Fig. S1). Hence, there were no differences in motor function that might have otherwise contributed to the differences in paw withdrawal threshold. Taking these findings together we conclude that while reduced expression of Beclin 1 does not affect basal pain threshold, male *Becn1^+/-^* are more hypersensitive to CFA than are the wild type controls, whereas female *Becn1^+/-^*mice were indistinguishable from wild type.

### Activating Beclin 1 reverses mechanical hypersensitivity in males

As inflammatory pain hypersensitivity was increased only in male *Becn1^+/-^*mice, we questioned whether activating Beclin 1 may suppress pain hypersensitivity in a sex-dependent manner. Beclin 1 is suppressed by endogenous inhibitors, including Golgi-associated plant pathogenesis related protein 1 (GAPR1), which binds to Beclin 1 preventing it from driving autophagy (*14*). This inhibitory interaction can be disrupted with an exogenous peptide that mimics the region in Beclin 1 which binds GAPR1. Linking this peptide to the HIV tat protein transduction domain allows the resultant peptide (tat-beclin 1) to be a cell-permeant activator of Beclin 1 (*14*). We tested the activity of tat-beclin 1 in the spinal dorsal horn by applying this peptide, or one in which the Beclin 1 sequence was rearranged (tat-scrambled), to dorsal horn neurons in primary culture (Fig. 2A). As a read out for Beclin 1 activity, we used the protein expression level of the autophagy substrate p62, which is inversely related to the activity of Beclin 1 and commonly used for measuring autophagy (*15*). We found that applying tat-beclin 1, but not tat-scrambled, reduced level of p62 (Fig. 2A), confirming activation of Beclin 1 by tat-beclin 1.

**Fig. 2.**
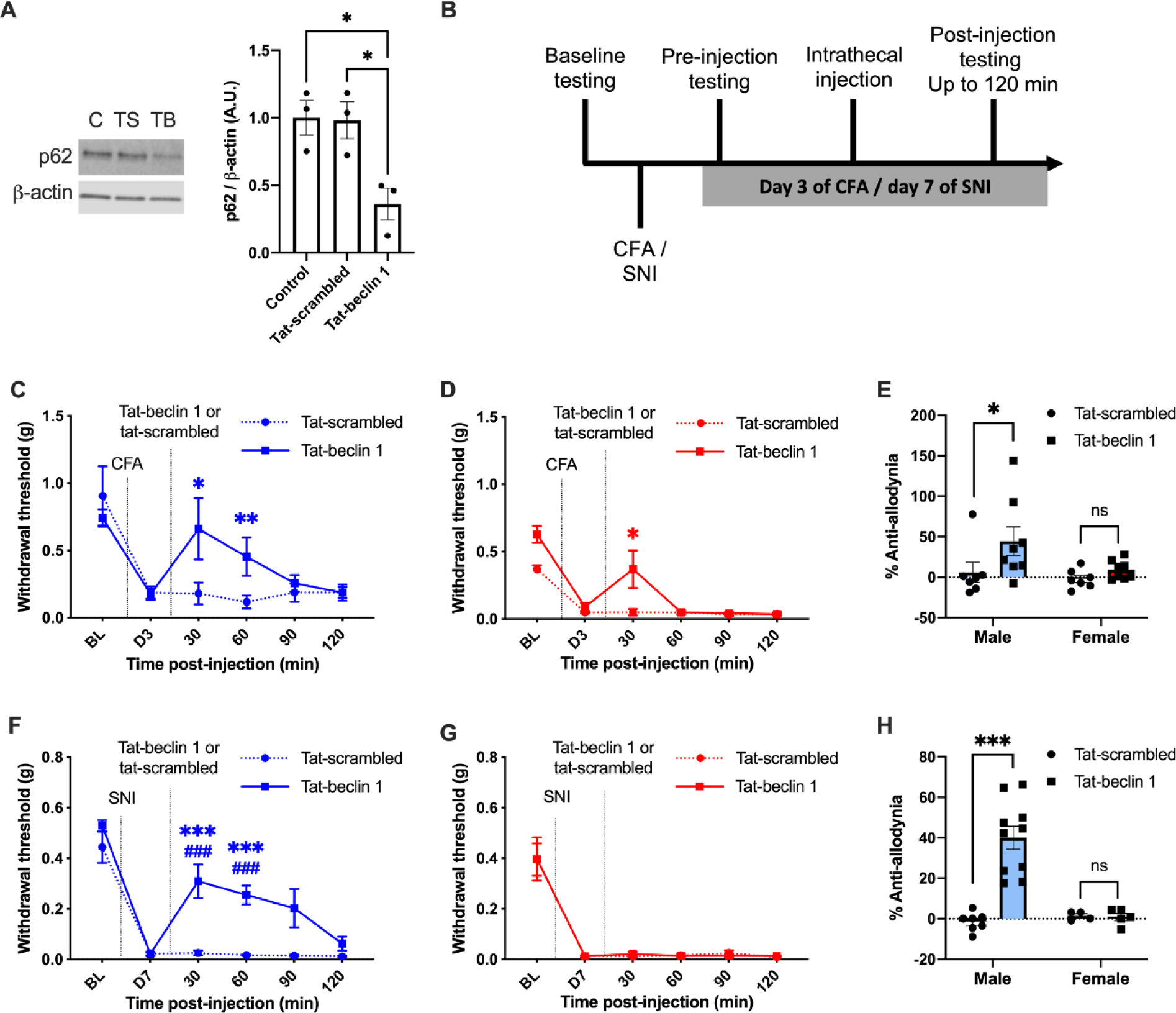
Beclin 1-activating peptide reverses inflammation- and nerve injury-induced mechanical hypersensitivity in males. **(A)** Protein expression of p62 in primary rat dorsal horn cultures treated with 120 µg/mL tat-beclin 1 (TB) or tat-scrambled (TS) for 2 hours. n=3; *p<0.05, one-way anova with Tukey post-hoc. **(B)** Experimental design timeline for intrathecal tat-beclin 1 experiments in Fig. 2C-H. **(C-D)** Mechanical PWT at baseline (BL), post-operative day 3 (D3) of CFA, followed by intrathecal injection of 7.5 µg tat-beclin 1 or tat-scrambled, in male (blue) and female (red) C57Bl/6 mice. **(E)** Anti-allodynia percentage of i.t. tat-beclin 1 or tat-scrambled in CFA quantified from Fig. 2C-D. CFA males tat-scrambled n=7, CFA males tat-beclin 1 n=8; CFA females tat-scrambled n=7, CFA females tat-beclin 1 n=8. **(F-G)** Mechanical PWT at baseline (BL), post-operative day 7 (D7) of SNI, followed by intrathecal injection of 7.5 µg tat-beclin 1 or tat-scrambled, in male (blue) and female (red) C57Bl/6 mice. **(H)** Anti-allodynia percentage of i.t. tat-beclin 1 or tat-scrambled in SNI quantified from Fig. 2F-G. SNI males tat-scrambled n=7, SNI males tat-beclin 1 n=10; SNI females tat-scrambled n=4, SNI females tat-beclin 1 n=5. *p<0.05, **p<0.01, ***p<0.001; *Drug vs. vehicle, Mann-Whitney U test; ^#^Timepoint vs. pre-injection D7 within group, Friedman test with Dunn’s post-hoc.

To investigate whether activating Beclin 1 affects inflammatory pain hypersensitivity, we administered tat-beclin 1 or tat-scrambled intrathecally in wild type mice that had developed hypersensitivity 3 days after intraplantar injection of CFA (Fig. 2B-E). In male mice, we found that administering tat-beclin 1 caused a statistically significant increase in PWT 30 and 60 minutes after injection compared to tat-scrambled, which had no effect on PWT, indicating reversal of the hypersensitivity induced by CFA (Fig. 2C). Moreover, tat-beclin 1 caused significant anti-allodynia, as measured by area under the curve (see Methods), over the two hours post-injection in the males (Fig. 2E). On the other hand, in female mice we found that tat-beclin 1 increased PWT at 30 minutes following intrathecal injection (Fig. 2D) and there was no overall anti-allodynic effect of this peptide (Fig. 2E).

To determine whether tat-beclin 1 affects neuropathic, as well as inflammatory, pain hypersensitivity, we tested the peptides in wild type mice using the spared nerve injury model (SNI), a widely used model of peripheral neuropathic pain (*16*). Tat-beclin 1 or tat-scrambled was administered intrathecally 7 days after nerve injury; all mice tested with the peptides had SNI-induced mechanical hypersensitivity (Fig. 2F-H). The PWT of male mice increased at 30 and 60 min after injecting tat-beclin 1 compared to tat-scrambled (Fig. 2F), and tat-beclin 1 produced significant anti-allodynia (Fig. 2H). By contrast, in female mice intrathecal injection of tat-beclin 1 did not affect PWT (Fig. 2G) nor did it cause anti-allodynia (Fig. 2H).

Taken together, these findings indicate that the Beclin 1 activating peptide tat-beclin 1 acutely reverses inflammatory and neuropathic pain hypersensitivity in male mice. In female mice, tat-beclin 1 had no effect on neuropathic pain hypersensitivity and an effect on inflammatory pain hypersensitivity that was less than that in male mice. Thus, we conclude that this activator of Beclin 1 causes a sex-dependent reversal of pain hypersensitivity in mice. Moreover, as reducing Beclin 1 expression exacerbates CFA-induced hypersensitivity (Fig. 1B), we suggest a model whereby, in males but not in females, pain hypersensitivity depends upon suppressing Beclin 1.

### Disrupting Beclin 1-Bcl2 interaction does not alter mechanical hypersensitivity

To determine whether the analgesic effects of tat-beclin 1 generalize to other activators of Beclin 1, we investigated the inhibition of Beclin 1 by binding of Bcl2 (*17*). Beclin 1 inhibition by Bcl2 depends upon phenylalanine at residue 121 in Beclin 1: mutating amino acid 121 on Beclin 1 from phenylalanine to alanine (F121A) reduces Bcl2 binding and increases autophagy (*18, 19*). Thus, we investigated whether the F121A mutation in mice affects basal nociception or hypersensitivity after inflammation or nerve injury. We found that baseline mechanical sensitivity in mice homozygously expressing alanine at residue 121 (*Becn1^F121A^*) was not different from that in wild type (*Becn1^+/+^*) mice in either males or females (Fig. 3). Following intraplantar injection of CFA, the reduction in PWT in *Becn1^F121A^* mice was indistinguishable from that in wild type in either sex over the 14-day period post-injection (Fig. 3A-D). Moreover, after SNI, allodynia in *Becn1^F121A^* was not different from that in wild type mice of either sex (Fig. 3E-H).

**Fig. 3.**
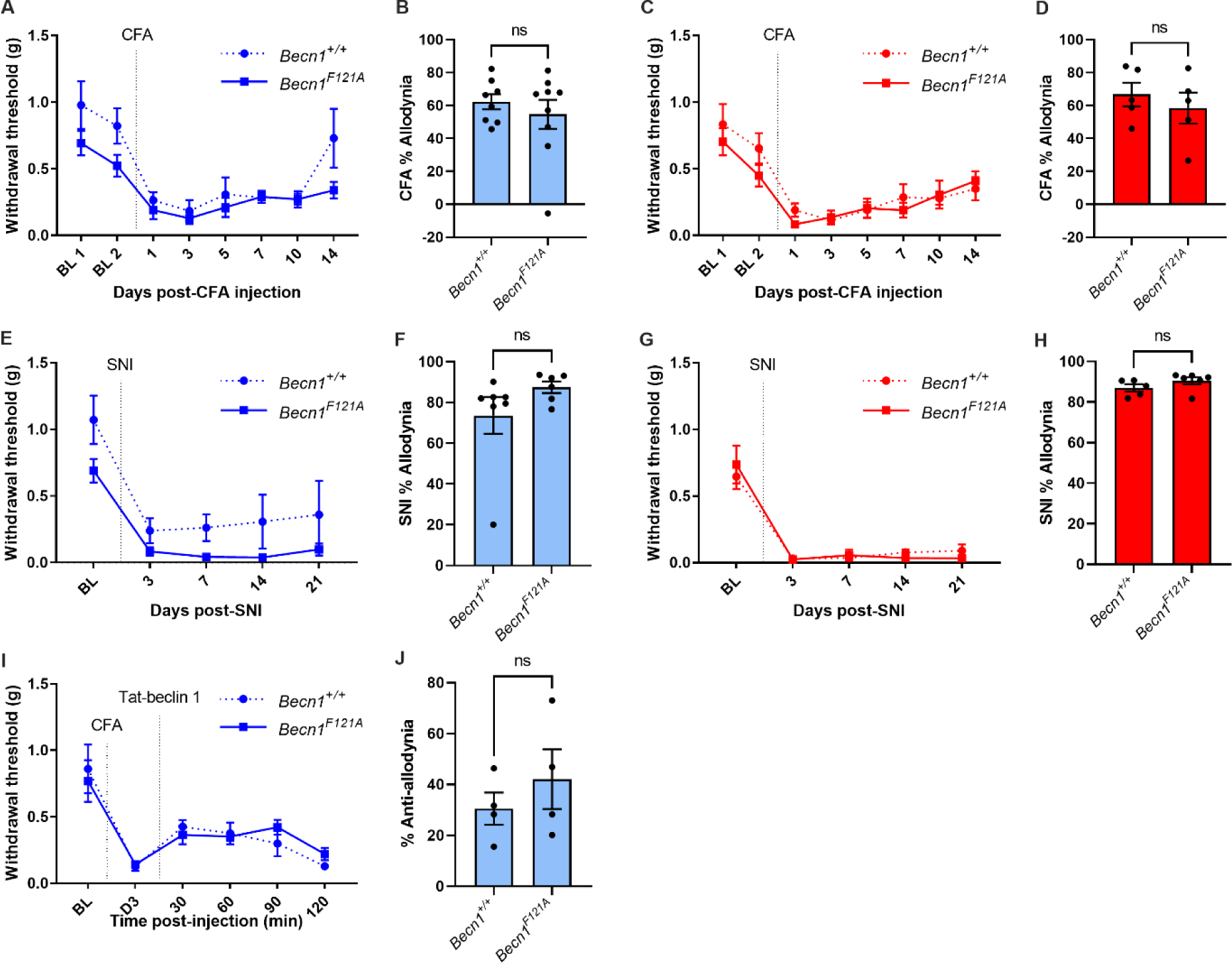
*Becn1^F121A^* mutation does not affect mechanical hypersensitivity. **(A-D)** Mechanical PWT of *Becn1^+/+^* (wild type) or *Becn1^F121A^* (homozygous mutants) male (blue) and female (red) mice at two baseline (BL) measurements and multiple days following intraplantar CFA injection, and expressed as overall percent allodynia. CFA males *Becn1^+/+^* n=8, CFA males *Becn1^F121A^* n=9; CFA females both genotypes: n=5. **(E-H)** Mechanical PWT of *Becn1^+/+^*or *Becn1^F121A^* male and female mice at BL and multiple days following SNI, and quantified as overall percent allodynia. SNI males *Becn1^+/+^* n=7, SNI males *Becn1^F121A^* n=6; SNI females *Becn1^+/+^* n=5, SNI females *Becn1^F121A^* n=6. **(I-J)** Mechanical paw withdrawal thresholds at BL, D3 of CFA, followed by intrathecal injection of tat-beclin 1 (7.5 µg) in *Becn1^+/+^* or *Becn1^F121A^* males, and expressed as percent anti-allodynia. n=4. *Becn1^+/+^* vs *Becn1^F121A^*, Mann-Whitney U test.

From the lack of effect of the F121A mutation on inflammatory or neuropathic pain hypersensitivity, we wondered whether other mechanisms regulating Beclin 1 activity remain important in regulating mechanical hypersensitivity despite the loss of Bcl2 inhibition. To explore this possibility, we tested whether tat-beclin 1 affects CFA-induced mechanical hypersensitivity in *Becn1^F121A^* mice. We found that administrating tat-beclin 1 intrathecally on day 3 post-CFA increased the PWT to the same extent in both *Becn1^F121A^* and wild type male mice (Fig. 3I-J). From these findings, we conclude that not all endogenous Beclin 1 inhibitory interactions participate in controlling pain hypersensitivity induced by CFA or SNI.

### BDNF-mediated mechanical hypersensitivity is dependent on Beclin 1

Sexual dimorphism in spinal pain signaling mechanisms is increasingly evident (*20*). A common mediator of inflammatory and neuropathic pain hypersensitivity is brain-derived neurotrophic factor (BDNF) (*21–23*), which is critical in males but not females (*24*). As here we found male-specific effects of manipulating Beclin 1, we tested whether the enhanced hypersensitivity of male *Becn1^+/-^* mice is dependent upon BDNF (Fig. 4A-C). In wild type males, we found that mechanical hypersensitivity induced by intraplantar CFA was reversed by intrathecally administering Y1036, a well-known inhibitor of BDNF signaling (Fig. 4A, C). By contrast, Y1036 had no effect on PWT in *Becn1^+/-^* male mice (Fig. 4A, C). In females, administering Y1036 had no effect on CFA-induced hypersensitivity in either wild type or *Becn1^+/-^*mice (Fig. 4B, C). The lack of effect of Y1036 in *Becn1^+/-^* males could not be attributed to a lack of responsivity to BDNF in these mice because we found that exogenously administering BDNF to naïve mice caused a decrease in the PWT in *Becn1^+/-^* males to the same extent as in wild type males (Fig. 4D). Thus, in *Becn1^+/-^* mice BDNF is not necessary for CFA-induced pain hypersensitivity. However, BDNF, when delivered exogenously, is sufficient to evoke hypersensitivity in either wild type or *Becn1^+/-^* male mice.

**Fig. 4.**
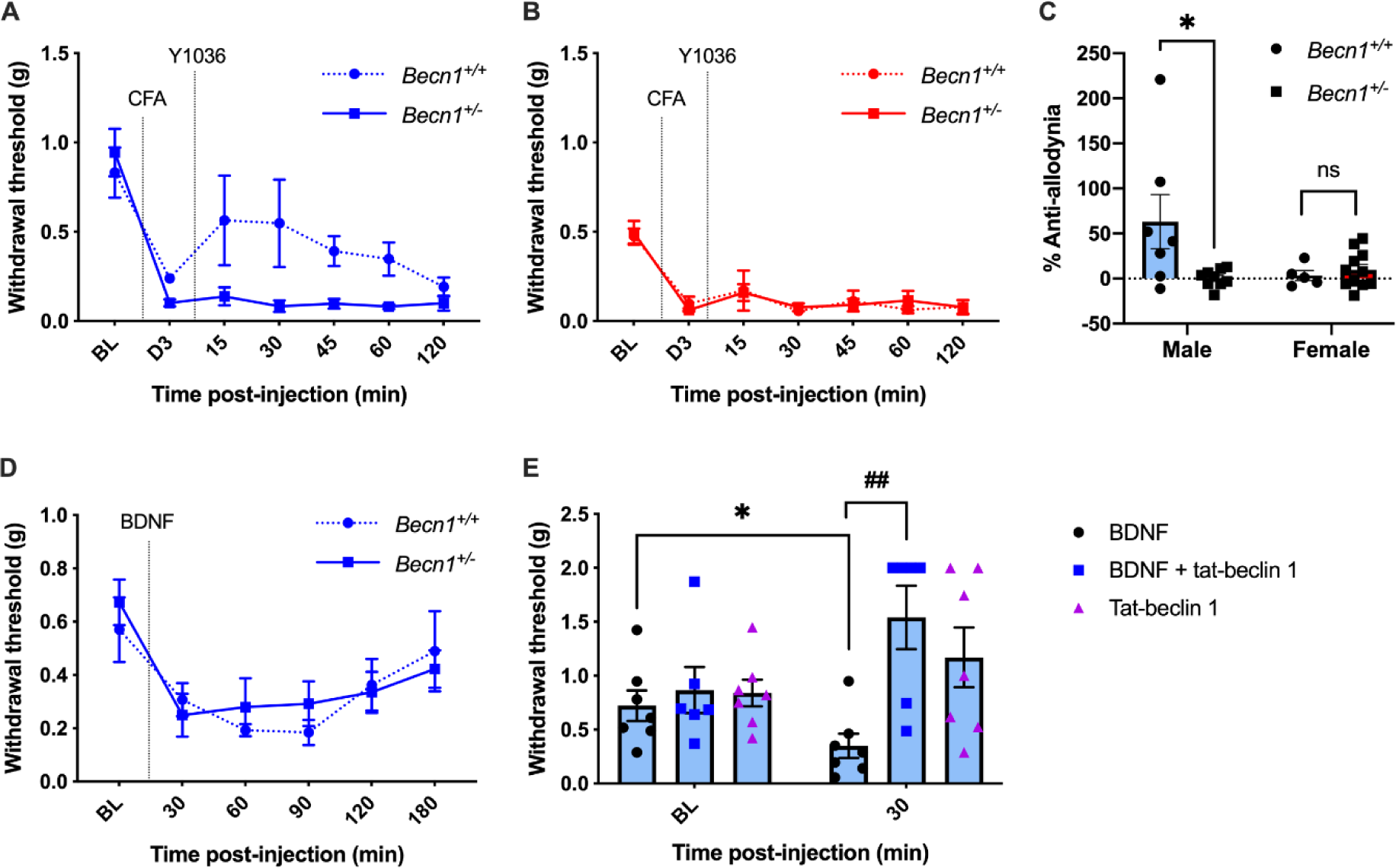
BDNF-mediated mechanical hypersensitivity is dependent on Beclin 1. **(A-C)** Mechanical PWT of *Becn1^+/+^* or *Becn1^+/-^* mice at baseline (BL), D3 of CFA, followed by intrathecal injection of the BDNF-inhibitor Y1036 (5 µg) in male (blue), and female (red). Also expressed as percent anti-allodynia for time course up to 60 min. Males *Becn1^+/+^*n=7, males *Becn1^+/-^* n=9; females *Becn1^+/+^* n=4, females *Becn1^+/-^* n=11. *p<0.05; *Mann-Whitney U test. **(D)** Mechanical PWT of *Becn1^+/+^* or *Becn1^+/-^* male mice at baseline, followed by intrathecal injection of BDNF (0.5 µg). *Becn1^+/+^*n=6, *Becn1^+/-^* n=10. **(E)** Mechanical PWT of naïve C57Bl/6 male mice following intrathecal injection of BDNF (0.5 µg), tat-beclin 1 (7.5 µg), or both. BDNF alone n=7, BDNF + tat-beclin 1 n=6, tat-beclin 1 alone n=7. *p<0.05, *Mann-Whitney U test; ^##^ p<0.01, ^#^Kruskal-Wallis test with Dunn’s post-hoc.

Our findings that, in wild type males, CFA-induced mechanical hypersensitivity depends upon BDNF (Fig. 4A) and is reversed by activating Beclin 1 (Fig. 2C) suggest that BDNF and Beclin 1 may act in a common signaling pathway in the spinal dorsal horn. This raises the possibility that BDNF may act upstream or downstream of Beclin 1. To investigate this possibility, we co-administered BDNF and tat-beclin 1 intrathecally in naïve, wild type male mice. We found that the reduction in PWT induced by administering BDNF alone was prevented by co-administering tat-beclin 1 (Fig. 4E). Importantly, tat-beclin 1 did not affect basal PWT without co-administering BDNF. Thus, our finding that activating Beclin 1 prevents hypersensitivity produced in naïve mice by exogenously delivered BDNF eliminates the possibility that in CFA-induced hypersensitivity Beclin 1 lies upstream of BDNF.

### Diminished Beclin 1 increases GluN2B expression and enhances NMDAR currents

Downstream of BDNF, signaling through its cognate receptor TrkB in dorsal horn neurons leads to upregulation of *N*-methyl-D-aspartate receptors (NMDARs) (*25, 26*). As NMDARs containing the GluN2B subunit are critical for driving pain hypersensitivity (*25–28*), we considered the possibility that reduction in Beclin 1, which exacerbates pain hypersensitivity in male but not female mice (see above), might increase the level of expression of NMDARs containing this subunit. In extracts from the dorsal spinal cord, we found that the level of GluN2B in male *Becn1^+/-^* mice was significantly greater than that in *Becn1^+/+^* mice (Fig. 5A), whereas the GluN2B level in female *Becn1^+/-^*mice was not different from that in *Becn1^+/+^* females (Fig. 5B). That *Becn1^+/-^* males have elevated GluN2B expression but no basal pain hypersensitivity in the absence of peripheral inflammation or nerve injury is consistent with the fact that NMDARs do not regulate PWT in naïve mice (*27, 29, 30*).

**Fig. 5.**
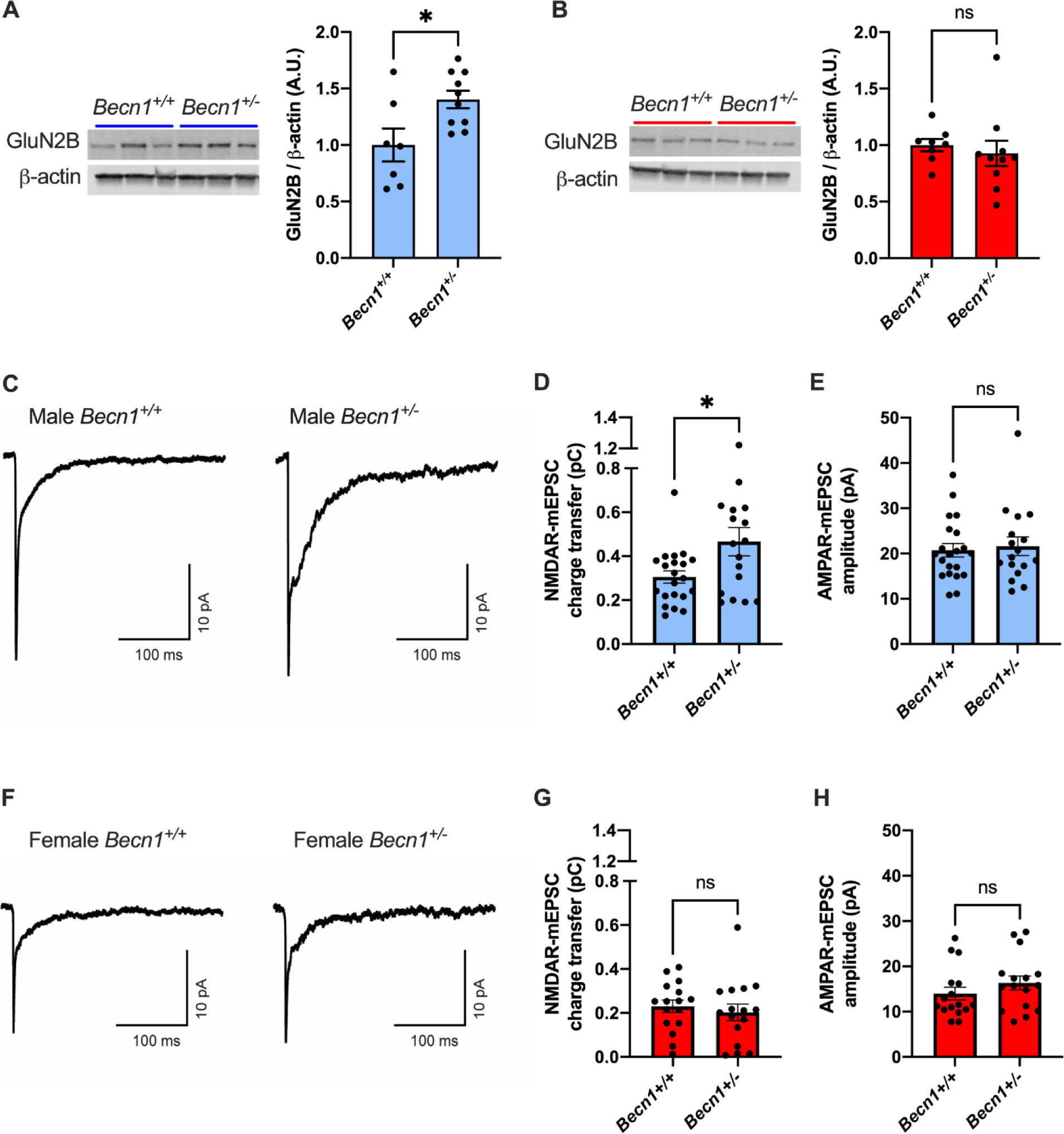
Monoallelic deletion of *Becn1* upregulates GluN2B in spinal dorsal horn neurons in males but not females. **(A-B)** Representative immunoblot images and quantification of GluN2B expression in spinal dorsal horn tissue from naïve *Becn1^+/+^* or *Becn1^+/-^* male mice (blue) and female mice (red). Males *Becn1^+/+^* n=7, males *Becn1^+/-^* n=10; females *Becn1^+/+^* n=8, females *Becn1^+/-^* n=10. **(C)** Representative traces of average mEPSCs recorded at −70 mV from cultured dorsal horn neurons from *Becn1^+/+^*or *Becn1^+/-^* male mice. **(D)** Magnitude of NMDAR-mediated charge transfer of mEPSCs from *Becn1^+/+^* or *Becn1^+/-^* male mice. **(E)** Magnitude of AMPAR-mediated amplitude of mEPSCs from *Becn1^+/+^* or *Becn1^+/-^* males. Males *Becn1^+/+^* n=21 cells, males *Becn1^+/-^* n=17 cells. **(F)** Representative traces of average mEPSCs recorded at −70 mV from cultured dorsal horn neurons from *Becn1^+/+^* or *Becn1^+/-^* female mice. **(G)** Magnitude of NMDAR-mediated charge transfer of mEPSCs from *Becn1^+/+^* or *Becn1^+/-^* females. **(H)** Magnitude of AMPAR-mediated fast component amplitude of mEPSCs from *Becn1^+/+^* or *Becn1^+/-^* females. Females both genotypes n=16 cells. *p<0.05, student’s t-test.

The greater expression of GluN2B in *Becn1^+/-^* males raises the possibility of increased responses mediated by NMDARs at synapses of dorsal horn neurons. We therefore compared miniature excitatory post-synaptic currents (mEPSCs) in dorsal horn neurons from *Becn1^+/-^* mice with those from the wild type (Fig. 5C-E). In recordings from neurons from male mice, we found that the NMDAR component of mEPSCs in *Becn1^+/-^* neurons was significantly greater than that in wild type neurons (Fig. 5D). On the other hand, in neurons from female mice the NMDAR component of mEPSCs from *Becn1^+/-^*neurons was indistinguishable from that of the wild type (Fig. 5F-G). While the NMDAR component of mEPSCs was increased in *Becn1^+/-^* male neurons, the AMPA receptor mediated component of the mEPSCs did not differ between *Becn1^+/-^* and wild type neurons in males (Fig. 5E), or in females (Fig. 5H). Moreover, we observed no differences in the frequency of mEPSCs between neurons from *Becn1^+/-^* and wild type mice in either sex (Fig. S2). Thus, reduced Beclin 1 in dorsal horn neurons from males, but not from females, produces a selective increase in synaptic NMDAR currents without observable alteration in synaptic transmission or in probability of glutamate release.

### Beclin 1 protein expression is unchanged after CFA

Our findings together support the model that hypersensitivity in pathological pain is dependent on reduced function of Beclin 1. This could be produced by either a reduced level of Beclin 1 protein or reduced Beclin 1 activity. To address whether Beclin 1 expression changes, in wild type mice we compared Beclin 1 protein level in the spinal dorsal horn of naïve mice with that of mice 3 days after CFA. We found that Beclin 1 protein expression in the lumbar dorsal horn did not differ in mice that received intraplantar CFA compared to naïve mice, in either males or females (Fig. 6). As it has been reported that the level of Beclin 1 protein in the dorsal spinal cord is also unchanged after SNI (*31*), we infer that Beclin 1 activity, rather than protein level, is downregulated after peripheral inflammation or nerve injury.

**Fig 6.**
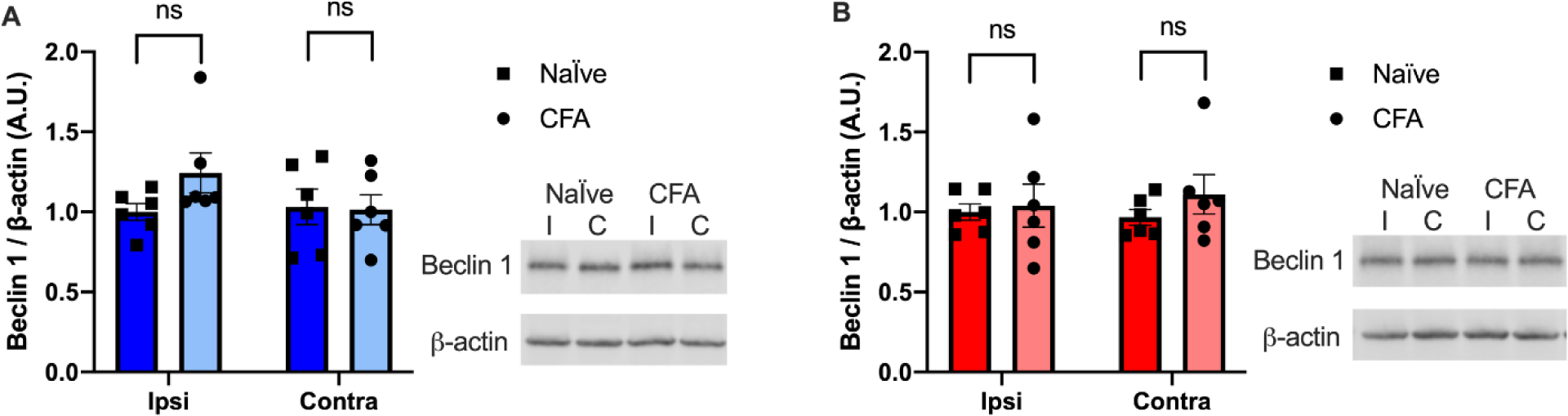
Beclin 1 expression in spinal dorsal horn does not change with intraplantar CFA. **(A-B)** Immunoblot of Beclin 1 protein expression in lumbar spinal dorsal horn ipsilateral (I) or contralateral (C) to the side of CFA injection. Tissue taken from naïve or day 3 of CFA C57Bl/6 males *(A)* and females *(B)*. n=6. Student’s t-test.

## DISCUSSION

Here, we provide converging lines of evidence for the involvement of Beclin 1 in pain hypersensitivity in male but not female mice. We found that inflammatory pain hypersensitivity, induced by CFA, was increased in male mutant mice with lowered expression of Beclin 1. Conversely, inflammatory pain hypersensitivity was reduced in wild type male mice, as was neuropathic pain hypersensitivity induced by SNI, by activating Beclin 1 with tat-beclin 1 administered at the level of the spinal cord. In contrast to hypersensitivity in the pain models, basal pain threshold was unaffected by either lowering expression or enhancing activity of Beclin 1, indicating it is selectively involved in pathological pain hypersensitivity. As the level of expression of Beclin 1 protein in the spinal cord was unchanged by CFA or SNI, the most parsimonious explanation for our findings taken together is that mechanical pain hypersensitivity after peripheral inflammation or nerve injury in males is dependent on inhibiting the activity of Beclin 1 in the spinal dorsal horn. In as much as inhibition of Beclin 1 reduces autophagy (*32*), we conclude that inhibiting autophagy in the spinal cord, which has been observed after SNI (*31, 33, 34*), mediates pathological pain hypersensitivity in males but not in females.

Inflammatory and neuropathic pain hypersensitivity share signaling by BDNF via activating its cognate receptor TrkB, as indicated by reversal of pain hypersensitivity by BDNF-TrkB inhibitors in wild type male rodents (*6, 21–24, 35, 36*). By contrast, we found that BDNF-TrkB signaling is dispensable for pain hypersensitivity in *Becn1^+/-^* mice. From this finding, together with our observation that activating Beclin 1 prevents hypersensitivity induced by exogenous BDNF, we infer that the inhibition of autophagy that mediates pain hypersensitivity is a consequence of BDNF-TrkB signaling. This inference is consistent with reports that BDNF can inhibit autophagy in hippocampal and cortical neurons via signaling through TrkB (*37, 38*). As a hub targeted by many autophagy modulatory pathways, Beclin 1 is known to be regulated by activating and inhibiting sites of phosphorylation through kinases including Unc-51 like autophagy-activating kinase 1 (ULK1), AMP-activated protein kinase (AMPK), mitogen-activated protein kinase-activated protein kinases 2 and 3 (MAPKAPK2/3), or AKT1 (*39*). Beclin 1 is also regulated by direct inhibitory interactions with proteins including GAPR1, Rubicon, and Bcl2 (*40*). While our findings with *Becn1^F121A^* mice eliminate the interaction with Bcl2 as necessary for regulating pain hypersensitivity (see Fig. 3), a major goal for future studies will be to determine the regulatory mechanisms by which BDNF-TrkB signaling suppresses autophagy to mediate pain hypersensitivity in the spinal dorsal horn.

In order to mediate pain hypersensitivity, autophagy needs to result in increased output of neurons in the nociceptive networks in the dorsal horn. A common mechanism leading to such increased output is enhancement of synaptic NMDARs, particularly those containing the GluN2B subunit (*25–27, 41, 42*). Here we found that reduced Beclin 1 expression led to elevated GluN2B expression in the dorsal horn and to enhanced NMDAR-mediated synaptic currents, in males only. NMDARs have been identified in proteomic profiling of autophagosomes from brain tissue (*43, 44*), demonstrating that NMDARs can be degraded via autophagy. Hence, a simple mechanism through which inhibition of autophagy may mediate pain hypersensitivity is by increasing NMDAR-mediated synaptic currents as a consequence of reducing autophagic degradation of GluN2B-containing NMDARs in spinal dorsal horn neurons. The male-specific increase in GluN2B and NMDAR currents in *Becn1^+/-^* mice is consistent with greater mechanical hypersensitivity after CFA in *Becn1^+/-^* males but not females. Also, that the elevated expression of GluN2B in *Becn1^+/-^* mice does not alter basal mechanical sensitivity is supported by the findings that while NMDARs are involved in pain hypersensitivity, these receptors do not participate in basal mechanosensation (*27, 29, 30*). The selective increase in NMDAR-mEPSCs but not AMPAR-mEPSCs in neurons from *Becn1^+/-^* mice phenocopies the selective increase in NMDAR-mEPSCs in lamina 1 dorsal horn neurons induced by peripheral nerve injury (*26*). Exogenously applied BDNF increases NMDAR-mESPCs in lamina 1 neurons from males but not females, not only in rodents but also in humans (*45*), which suggests cross-species conservation of the role of autophagy in mediating pathological pain hypersensitivity.

Taken together we propose that, in males, peripheral inflammation and nerve injury induces release of BDNF in the spinal dorsal horn; BDNF signaling inhibits Beclin 1, leading to enhanced NMDAR-mediated currents in dorsal horn neurons, resulting in pain hypersensitivity. We demonstrate that pharmacologic activation of Beclin 1 with tat-beclin 1 reverses pain hypersensitivity in males, identifying Beclin 1 as a novel therapeutic target and providing the basis for a new class of drugs for chronic pain.

## MATERIALS AND METHODS

### Animals

All animal experiments were approved by the Hospital for Sick Children’s Animal Care Committee and in accordance with the Canadian Council on Animal Care (CCAC) guidelines. *Becn1^fl/fl^* mice obtained from Martin Post’s lab (Hospital for Sick Children) were crossed with CMV-cre (Jackson) to generate whole body *Becn1^+/-^* mice. *Becn1^+/-^* CMV-Cre-positive mice were then crossed with wild type C57Bl/6J mice to generate *Becn1^+/-^* cre-negative mice. Certain experiments with *Becn1^+/-^* contained both cre-positive and cre-negative mice; cre expression caused no significant difference in outcomes measured. Genotyping for *Becn1^+/-^* was performed as described (*11*). *Becn1^F121A^* mice were obtained from Beth Levine’s lab (University of Texas Southwestern) and bred with wild type C57Bl/6J mice. Genotyping for *Becn1^F121A^* was performed as described (*18*).

### Intraplantar CFA

Under isoflurane/oxygen anesthesia, mice received unilateral injection of CFA (Sigma) (30% CFA in 20 µL saline emulsion) into the plantar surface of the left hind paw.

### Spared nerve injury (SNI)

Mice received unilateral SNI as previously described (*16*). Briefly, mice under isoflurane/oxygen anesthesia received an incision on the left thigh and blunt dissection of the biceps femoris muscle to expose the sciatic nerve. The tibial and common peroneal nerves were ligated with 6-0 vicryl sutures above the branch, then transected. The sural nerve was left intact. The muscle and skin incisions were sutured closed.

### Rotarod

Mice were placed on the Rotarod apparatus (Panlab) moving in the direction opposite of the rotarod rotation. Rotarod was set at initial speed of 4 rpm with constant acceleration to maximum 40 rpm within 30 seconds. Mice trained on the rotarod 3 times before testing trials. Latency to fall off the rotarod was reported as an average of 3 testing trials. Experimenters were blinded to genotype. Mice age ranged from 8-19 weeks old.

### Von Frey behaviour testing

Mice were placed in plastic cubicles on a perforated metal platform and habituated for at least 60 min before testing. To assess mechanical sensitivity, nylon monofilaments (Stoelting Touch Test) were applied to the mid-plantar hind paw for CFA experiments or the lateral aspect of the hindpaw for SNI experiments. The 50% mechanical withdrawal threshold was calculated using the up-down method of Dixon (*46*) from the average of two measurements taken at least 15 min apart. Post-intrathecal injection withdrawal thresholds were taken from single measurements. Experimenters were blinded to genotype and drug treatment. Mice age ranged from 6-17 weeks old. One *Becn1^+/+^*mouse was excluded from Fig. 3G-H because of hyposensitivity after SNI.

Quantification of percent anti-allodynia by drugs was performed using area under the PWT-time curve (using trapezoidal method) over the post-intrathecal injection testing period, then reported as a percentage of maximum possible anti-allodynia (a hypothetical area under the PWT-time curve where the drug brings PWT from the post-injury (CFA D3 or SNI D7) back to the baseline PWT of the individual mouse at each post-injection timepoint), as previously described (*24*).

Percent allodynia for a 14 or 21 day time course was calculated as area over the curve, by subtracting area under the curve of the PWT-time curve (using trapezoid method) from the hypothetical maximum area, with the individual’s average baseline PWT as maximum PWT. Then expressed as a percentage of the maximum possible allodynia.

### Drug administration

Intrathecal injections were performed using a 30 gauge, ½ inch needle between the L5 and L6 vertebrae of the mouse under light restraint. Tat-beclin 1 (YGRKKRRQRRRGGTNVFNATFEIWHDGEFGT) (7.5 µg, Genscript) or tat-scrambled (YGRKKRRQRRRGGVGNDFFINHETTGFATEW) (7.5 µg, Genscript) dissolved in 5 µL PBS. BDNF (0.5 µg, Alamone) was dissolved in 10 µL PBS. Co-injection of BDNF and tat-beclin 1 were administered at the same doses together, in a final volume of 10 µL. Y1036 (5 µg, Calbiochem) was dissolved in 5 µL of 2% DMSO in water.

### Primary dorsal horn cultures

Cultures of dorsal spinal cord were prepared from fetal Wistar rats (Charles River) on embryonic day 17 (E17) or *Becn1^+/-^* or *Becn1^+/+^* mice on E15. The fetuses were removed from time-pregnant females immediately after euthanasia by ether (rat) or isoflurane (mouse) followed by cervical dislocation. Upon removal the fetuses were transferred to chilled sterile Hanks’ buffered salt solution and were then euthanized by decapitation. The entire spinal cord was removed by an anterior approach and was placed dorsal side up in iced Hanks’ solution. The meninges and dorsal root ganglia were removed. The dissection procedure used to isolate the dorsal spinal cord was similar the “open-book” technique as previously described (*47*). The cord was opened by incising it through the dorsal commissure to the central canal along the entire length. The opened cord was turned over and pinned onto a dish coated with Sylgard (Dow-Corning). The dorsal half of the cord was removed by cutting along the lateral funiculus on each side. The dorsal halves of cords were pooled and minced. For rat dorsal horn, the cords were incubated for 25 min at 37° C in Hanks’ solution with trypsin (0.25%). The trypsin was inactivated by rinsing the dorsal cords in Neurobasal media with 1% fetal bovine serum, B27 and L-Glutamine. The tissue was then mechanically dissociated by trituration. The cells were plated onto 12 mm glass coverslips, coated with Poly D-Lysine (Sigma). Cells were maintained in Neurobasal media (Gibco) supplemented with 1% fetal bovine serum (FBS), L-Glutamine and B27 for 3-4 days. After 3-4 days, media was switched to Neurobasal media with supplements but no FBS. Cells were used at 13-18 days in culture.

### Electrophysiology

Mouse dorsal horn neurons, cultured on 10-mm coverslips for 13-18 days, were transferred to 35-mm petri dishes, and then whole-cell patch-clamp recordings were made at room temperature (22 °C). A MultiClamp 700B microelectrode amplifier (Molecular Devices) and an Axon Digidata 1440A acquisition system (Molecular Devices) were used for electrophysiological recordings. Electrical signals were filtered at 2 kHz and sampled at 10 kHz. A P-87 pipette puller (Sutter Instrument Co.) was used for pulling recording electrodes with micropipettes (World Precision Instruments). The electrodes had resistance of 3-5 MΩ, and were filled with the intracellular recording solution composed of (in mM): 137 CsF, 1.5 CsCl, 10 BAPTA, 4 Mg-ATP and 10 HEPES. The pH of the intracellular solution was adjusted to 7.20 with CsOH. The extracellular recording solution in 35-mm dishes contained (in mM): 140 NaCl, 5.4 KCl, 15 HEPES, 25 glucose, 0.0005 tetrodotoxin, 1.3 CaCl_2_, 0.01 bicuculline, 0.001 strychnine, and 0.001 glycine, and pH was adjusted to 7.35 with NaOH. Miniature excitatory postsynaptic currents (mEPSCs) were recorded at the membrane potentials of −70 mV under voltage-clamp condition. The experimenter was blinded to genotype during the recordings and analysis. Recording data were analyzed offline using Clampfit 10.7 software (Molecular Devices) and Mini Analysis Program 6.0 (Synaptosoft Inc). AMPA receptor- and NMDA receptor-mediated mEPSC components were determined as reported previously (*48*). The AMPA receptor-mEPSC component was measured at peak amplitude of averaged mEPSCs, and the NMDAR receptor-mEPSC component was determined through the receptor charge transfer by integrating the area from 25 ms to 225 ms after the peak value.

### Immunoblot

Lumbar spinal dorsal horn, or specifically L4/5 region of the lumbar dorsal horn for CFA experiments, were isolated from mice and tissues were frozen in −80 °C until processed. Tissues or cultured cells were homogenized in radioimmunoprecipitation assay (RIPA) buffer (50 mM Tris-HCl, 150 mM NaCl, 2mM EDTA, 0.1% SDS, 1% NP-40, 0.5% deoxycholate, protease inhibitor cocktail, and phosphatase inhibitor cocktails), incubated in RIPA buffer at 4 °C for at least 30 min, then centrifuged at 14 000 rpm for 15 min at 4 °C. Proteins in the supernatant were boiled in laemmli buffer with DTT, then equal amounts of protein per sample were separated by SDS-PAGE (Bio-Rad) and transferred to PVDF membranes (Immobilon). Membranes were blocked with Intercept blocking buffer (LI-COR) for 1 hr at room temperature, then incubated overnight at 4 °C in primary antibodies: mouse anti-Beclin 1 (Abcam), mouse anti-GluN2B (BD Transductions), mouse anti-β-actin (Sigma), or rabbit anti-β-actin (Cell Signaling). Membranes were washed with Tris-buffered saline with 0.1% Tween-20, then incubated in fluorescent secondary antibodies (LI-COR) for 1 hr at room temperature, washed, then imaged with LI-COR Odessey FC and quantified with Image Studio Lite 5.2 (LI-COR).

### Statistical analysis

Data are presented as means ± SEM. Statistical analysis was performed using GraphPad Prism 9, with statistical significance set as p < 0.05. Statistical tests are described in figure legends for each corresponding experiment.

## ACKNOWLEDGEMENTS

The authors sincerely thank Yongqian Wang for technical assistance, Dr. Ameet Sengar for scientific discussion, and Dr. Beth Levine and Dr. Martin Post for providing mouse strains. This research was supported by funding from the Canadian Institutes for Health Research (CIHR) to MWS (FDN-154336). THT was supported by scholarships from CIHR, Ontario Graduate Scholarships, Ontario Women’s Health Scholars Award, and Restracomp scholarship from the Hospital for Sick Children.

## SUPPLEMENTARY MATERIALS

**Fig. S1.**
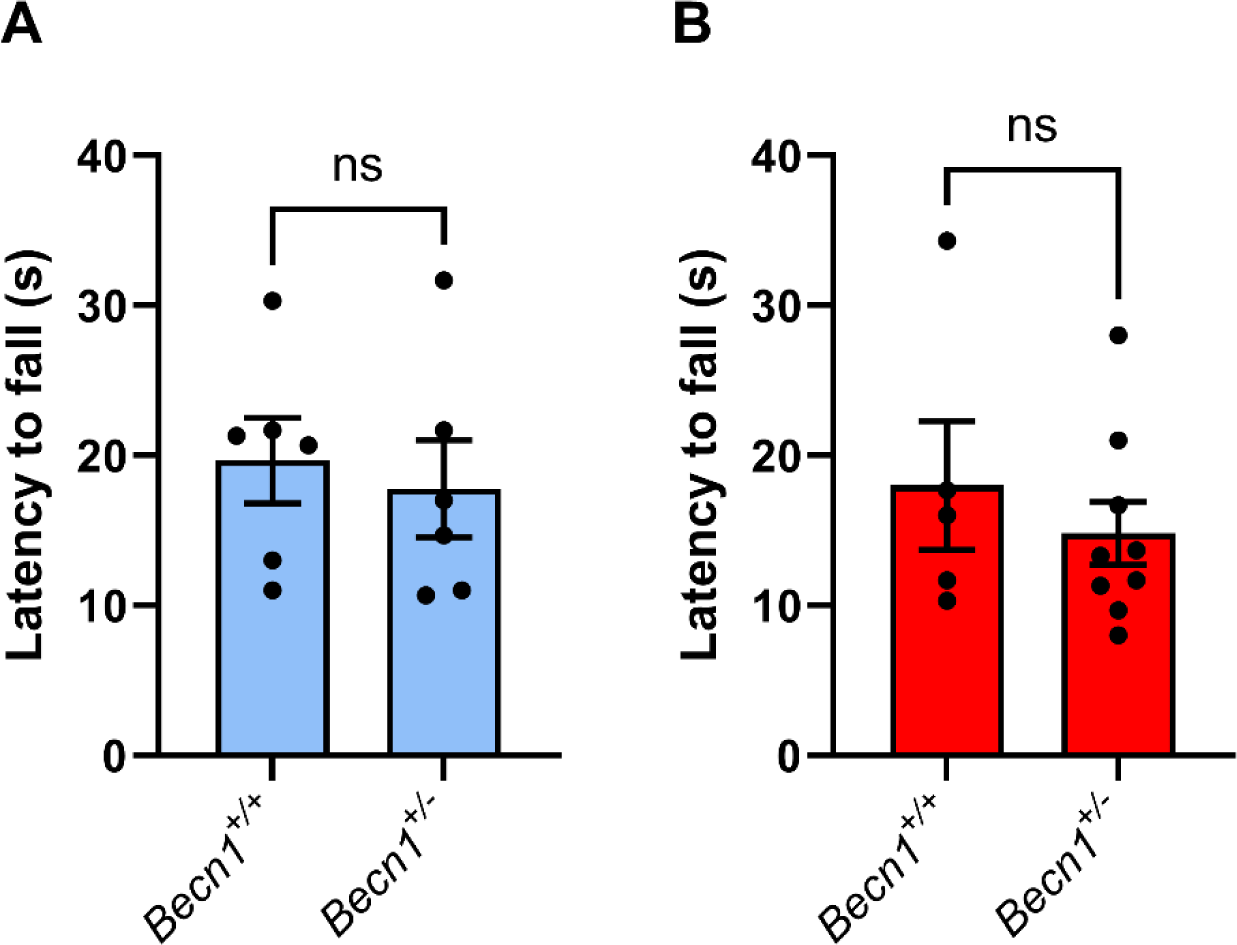
Monoallelic deletion of *Becn1* does not affect performance on rotarod test. Accelerating rotarod test latency to fall of *Becn1^+/+^* or *Becn1^+/-^* males *(A)* and females *(B)*. Males both genotypes: n=6; females *Becn1^+/+^* n=5, females *Becn1^+/-^* n=9. Not significant (ns), student’s t-test.

**Fig. S2.**
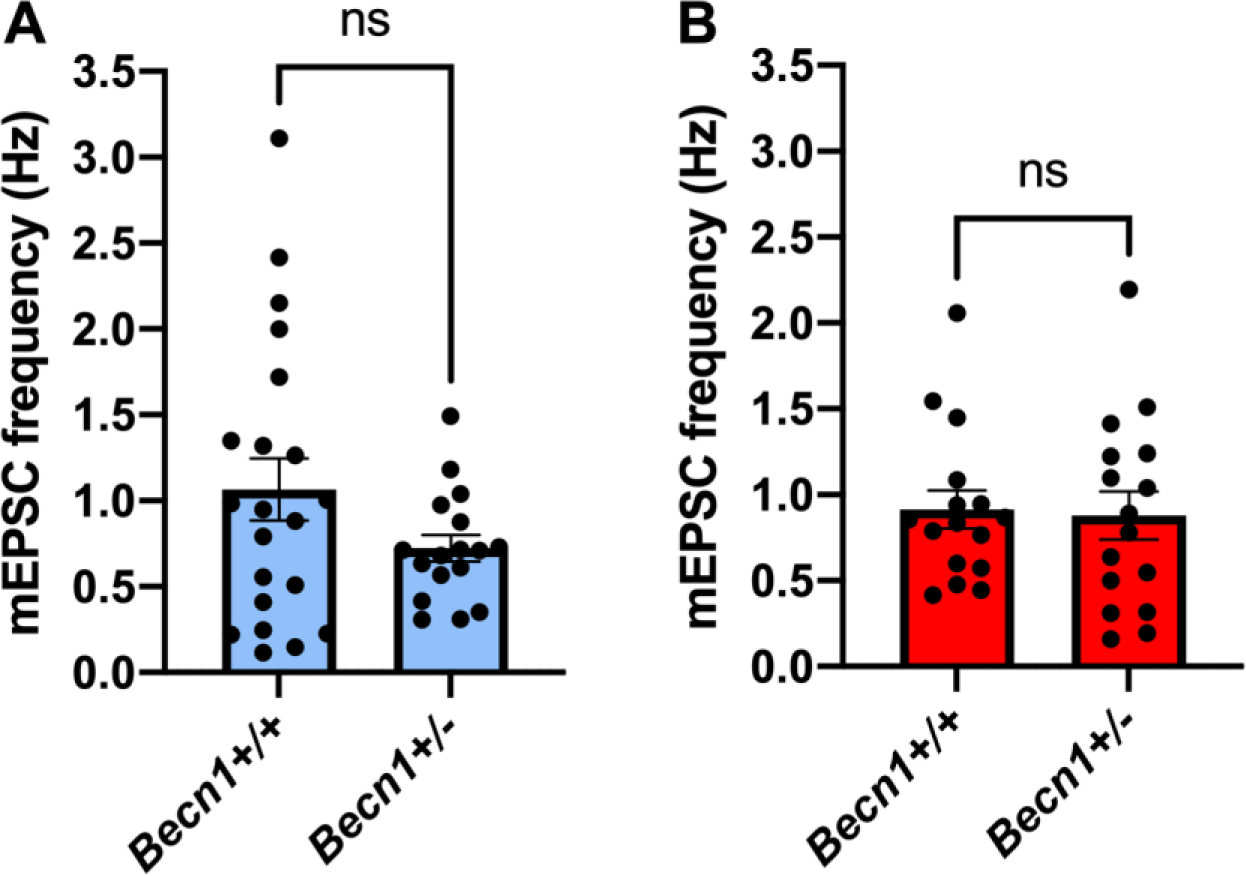
Monoallelic deletion of *Becn1* does not affect frequency of mEPSCs. Frequency of mEPSCs recorded from cultured spinal dorsal horn neurons from *Becn1^+/+^* or *Becn1^+/-^* males *(A)* and females *(B)*. Males *Becn1^+/+^* n=21 cells, males *Becn1^+/-^*n=17 cells; females both genotypes n=16 cells. Not significant (ns), student’s t-test.

